# Assessing the potential for genome-assisted breeding in red perilla using quantitative trait locus analysis and genomic prediction

**DOI:** 10.1101/2023.10.15.560967

**Authors:** Sei Kinoshita, Kengo Sakurai, Kosuke Hamazaki, Takahiro Tsusaka, Miki Sakurai, Terue Kurosawa, Youichi Aoki, Kenta Shirasawa, Sachiko Isobe, Hiroyoshi Iwata

## Abstract

Red perilla is an important medicinal plant used in Kampo medicine, and the development of elite varieties of this species is urgently necessary. Medicinal compounds are generally considered target traits in medicinal plant breeding; however, selection based on the phenotypes of the compounds (i.e., conventional selection) is expensive and time-consuming. Here, we proposed genomic selection (GS) and marker-assisted selection (MAS) as suitable selection methods for medicinal plants, and evaluated the effectiveness of GS and MAS in red perilla breeding. Three breeding populations generated from crosses between one red and three green perilla genotypes were used to elucidate the genetic mechanisms underlying the production of major medicinal compounds using quantitative trait locus analysis and to evaluate the accuracy of genomic prediction (GP). We found that GP had sufficiently high accuracy for all traits, confirming that GS was an effective method for perilla breeding. Moreover, the three populations showed varying degrees of segregation, suggesting that use of these populations in breeding may yield simultaneous enhancements of different target traits. This study can contribute to research on the genetic mechanisms of the major medicinal compounds of red perilla as well as the breeding efficiency of this medicinal plant.

## 1. Introduction

*Perilla frutescens*, an annual herb of the Labiatae family native to China, is widely cultivated in China, Japan, Korea, and other Asian countries [1]. *P. frutescens* has long been used as food and an ingredient in traditional (Kampo) medicine. *Perilla frutescens* can be categorized into two types based on morphological differences in its leaves: red perilla (‘aka-jiso’), which has dark red or purple leaves and stems due to the presence of anthocyanin, and green perilla (‘ao-jiso’), which only contains a small amount of anthocyanin and has green leaves [2,3]. Red perilla is used as an ingredient in Kampo medicines, often to ease stomach, cold, and anxiety, and is blended in several formulations such as Hangekobokuto and Kososan. The main medicinal compounds in perilla are perillaldehyde (PA), rosmarinic acid (RA), and anthocyanin (ANT) [4]. PA is an odorant with antidepressant, anticancer, and antibacterial properties; RA is a phenol with anti-inflammatory and antioxidant properties and is more abundant in green perilla; and ANT is a red pigment with antioxidant properties [2,3,5]. Owing to its benefits, red perilla has attracted attention as an ingredient for Kampo medicine both in Asian countries and globally in recent years [6]. The development of varieties with stable medicinal compounds is necessary to meet the increasing demand for red perilla. Although extensive research has been conducted on the bioactivity, metabolic pathways, and efficacy of the medicinal compounds of red perilla, no research has been conducted on the enhancement of these medicinal compounds via selective breeding.

The quality of perilla plants used in Kampo medicine is determined by the concentrations of their major medicinal compounds, PA, RA, and ANT. Therefore, these compounds are considered target traits when breeding red perilla as a medicinal plant. Conventional phenotype-selection-based breeding is not an efficient breeding method for medicinal plants, as it necessitates the quantification of medicinal compounds in every generation and individual, which is both expensive and time-consuming. Compound analysis involves steps such as sampling from the plant, drying, sample grinding, reagent preparation, compound extraction, quantification using ultra high-performance liquid chromatography (UPLC), and organizing data. Even for experienced individuals, these steps take over 1 month to complete, during which time no other experiments can be conducted. Additionally, there are economic costs involved in terms of field management and labor. Genomic selection (GS) [7], an alternative approach wherein genomic prediction (GP) is used to predict phenotypic values from genome-wide marker information, is also suitable for improving medicinal traits. In GS, once a GP model is built, phenotypes can be predicted based solely on genomic information without the need to quantify medicinal compounds in every generation. If the target trait is controlled by a small number of quantitative trait loci (QTLs), marker-assisted selection (MAS), a method of selection using markers that are strongly linked to the QTLs that control the target trait, can also be used for selection using only the genotypes of the linked markers. Genomic information can be obtained during the early growth stages of a plant, eliminating the need for cultivation in the field and accelerating the breeding cycle. With the development of next-generation sequencing (NGS) technologies, obtaining genomic information has become easier and inexpensive. Therefore, GS and MAS are better suited than phenotypic selection for breeding red perilla.

GS was first proposed by Meuwissen et al. in 2001 and has been implemented successfully in dairy cattle breeding [7,8]. In plant breeding, GS has been applied in major crops such as maize and rice as well as other plants [9–11]. Most previous studies using GS have focused on yield or weight as target traits; however, some studies have focused on nutrients and other active ingredients, such as vitamin E in sweet corn, carotenoid in maize, or various metabolites in oat [12–14]. GS focusing on the active ingredients is a crucial method in plant breeding. However, no studies have investigated GS in medicinal plants using medicinal compounds as target traits. Furthermore, in breeding to improve medicinal compounds, it is essential to target multiple compounds simultaneously; however, few such studies have been conducted. When several traits are improved simultaneously, it is important to consider both the correlation among the traits as well as the correlation between medicinal compounds and other important agronomic traits, such as yield and flowering date. Therefore, it is insufficient to focus only on the phenotypic correlations among these traits; genetic architecture and genetic correlations estimated by genetic relationship matrices should also be utilized to assess whether each trait could be improved independently.

As few elite cultivars of red perilla for medicinal use exist, it is difficult to improve multiple medicinal compounds using only one population generated through a single cross combination. In this study, to enhance multiple medicinal compounds simultaneously, we used three populations generated by crossing red and green perilla. When multiple populations with different genetic backgrounds are used, the genetic correlations among traits, heritability, and accuracies of GPs are likely to differ among populations. Therefore, it is necessary to evaluate the extent of variation in these values across populations and whether the results differ from those obtained when predictions are made using all the three populations together.

The present study had two main objectives: to evaluate the possibility of improving major medicinal compounds and other agronomic traits simultaneously in red perilla through genomic breeding, and to identify the genetic characteristics of each population arising from differences in the effects of alleles derived from different parents and different levels of segregation in later generations of the population. To achieve these objectives, we determined the genetic correlations among the traits mentioned above and conducted QTL analysis and GP using three populations with different genetic backgrounds. Based on the results of these analyses, the genetic architecture of each trait and the potential for breeding using GS and MAS are discussed. The results of this study provide valuable insights for studying the genetic mechanisms underlying the production of the major medicinal compounds of red perilla and will contribute substantially to red perilla breeding.

## 2. Materials and Methods

### 2.1. Plant Materials

The plant materials used in this study were the breeding populations of the F_3_ and F_4_ generations of two-way crosses of red perilla and green perilla. The breeding population for each generation consisted of three populations. The cross-parents of these populations were ‘SekihoS8,’ st27, st40, and st44. ‘SekihoS8’ is a representative variety of red perilla (*P. frutescens* var. *crispa f. purpurea*) owned by TSUMURA &CO., and st27, st40, and st44 are green perilla (*P. frutescens Britton* var. *crispa Decne*) owned by the National Agricultural and Food Research Organization. The cross combinations were ‘SekihoS8’ × st27 (S827), ‘SekihoS8’ × st40 (S840), and ‘SekihoS8’ × st44 (S844), hereafter referred to as S827, S840, and S844, respectively. The plants were grown in Ibaraki field, TSUMURA & CO., in 2021 and 2022.

The selection and breeding scheme used to derive the breeding populations are shown in Figure 1. The initial cross parents were as described above, and the breeding scheme was advanced by repeated self-pollination. In the F_2_ generation, 200 lines were grown from each of the three cross combinations. From these lines, 100 lines from each population were selected for self-pollination, and three seeds per line were collected and cultivated for the F_3_ generation. In the F_3_ generation, to maintain the genetic diversity of the population, 50 lines were selected from the lines that were grown in the field, and 50 other lines were selected from those not grown in the field for each population. Three individuals were cultivated as the next generation, F_4_, from seeds collected from each line.

**Figure 1.**
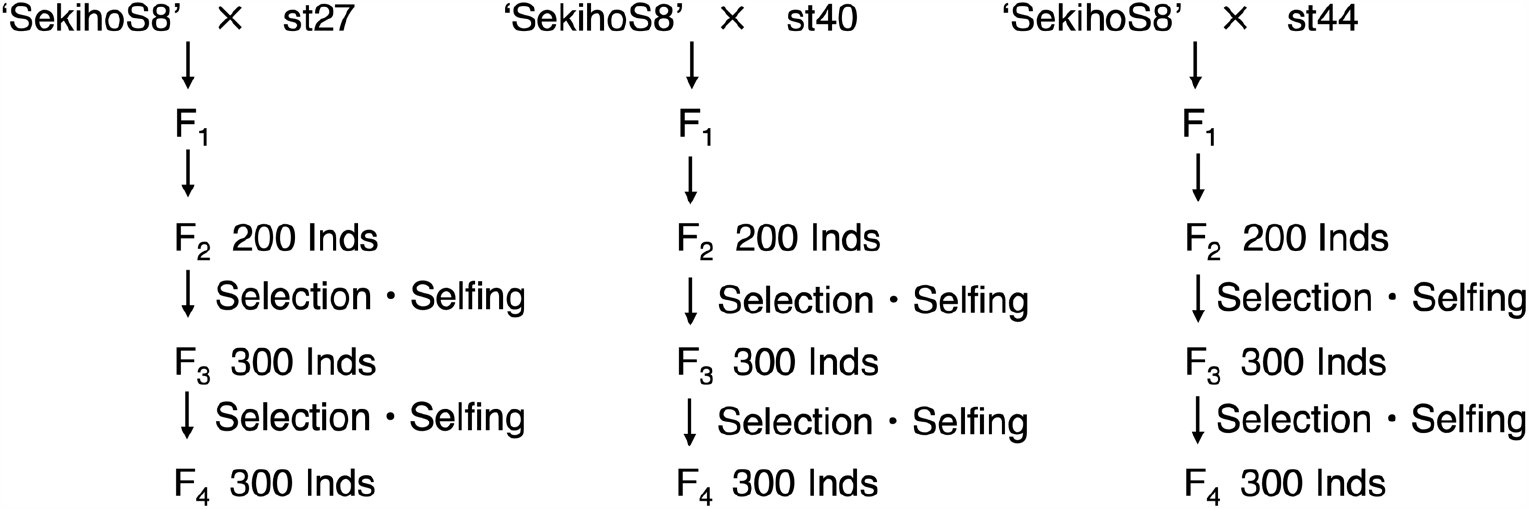
Selection and breeding scheme of the breeding populations used as plant parent materials in this study.

### 2.2. Phenotype Data

In the F_3_ population, the contents of PA, RA, and ANT were measured in plants grown in the experimental field. In the F_4_ population, PA, RA, yield, and flowering date were measured in plants grown in the experimental field.

Quantification of PA and RA was performed as reported previously [15,16]. To determine the PA content, 0.1 g of each plant was weighed, placed in 25 mL of methanol, shaken, and centrifuged. The supernatant was then separated and analyzed via UPLC at a wavelength of 234 nm using a UV detector. To determine the RA content, 0.05 g of each plant was weighed, placed in a 25 mL methanol/water (3:1) solution, shaken, and centrifuged, after which the supernatant was separated and analyzed via UPLC at a wavelength of 330 nm using a UV detector. The ANT was measured three times on July 14, July 29, and August 12, 2021. For each measurement, the ANT content was measured using three fully expanded leaves per plant with an anthocyanin content meter, and the average was recorded. The yield was measured three times on July 19, July 29, and August 8, 2022, and all three measurements were added for later analysis. For each yield measurement, plants were harvested at a height of 45 cm using a machine and the weight of the harvested plants was measured. The survey on flowering dates was conducted from August 31, 2022, to September 30, 2022. The number of days from April 25, the date of sowing in the cell trays, to the flowering date was used for data analysis.

### 2.3. Genotype Data

Genomic DNA was extracted from all the individuals in the F_3_ and F_4_ generations as described in Subsection 2.1, as well as on the four parental lines, ‘SekihoS8,’ st27, st40, and st44. Double-digest restriction-associated DNA sequencing [17] was performed using a DNBSeq-G400RS (MGI Tech Co. Ltd. Shenzhen, China) to examine genome-wide single nucleotide polymorphisms (SNPs). The 100 paired-end reads obtained were mapped to the reference sequence Hoko-3 (*P. frutescens cv*.) published by Tamura et al. [18] using Bowtie2 [19]. Variant calling with max-missing 0.9 was conducted, and low-quality SNPs were filtered out using VCFtools version. 0.1.16 [20], after which 2,282 SNPs were identified. SNPs with minor allele frequencies (MAF) <0.025 were excluded, and missing data were input using Beagle version. 5.1 [21], after which they were filtered again for an MAF < 0.025. Finally, 2,063 SNPs were identified. These SNPs were filtered with MAF < 0.01 for both each population and the total three populations. In the F_3_ generation, 1,964 SNPs remained in the entire population: 727 SNPs in S827, 1,432 SNPs in S840, and 579 SNPs in S844. In the F_4_ generation, 1,964 SNPs remained in the entire population: 862 SNPs in S827, 1,432 SNPs in S840, and 579 SNPs in S844. For later analysis, the score for SNPs of the same genotype as ‘SekihoS8’ was set to 0, 1 for heterozygous SNPs, and 2 for homozygous SNPs in the other parent.

### 2.4. Linkage Map Construction and QTL Analysis

The F_3_ generation was used to create a linkage map, and the 1,964 SNPs (S827: 725 SNPs, S840: 1,432 SNPs, and S844: 579 SNPs) described in the previous subsection were used as marker genotypes. The order of the markers was the same as that described in Subsection 2.3. Based on the expected ratio of 3:2:3 for marker classes at each locus, the Maximum Likelihood Method was used to estimate the recombination rates between markers for each population. Markers with recombination rates greater than 0.499 were excluded and the remaining markers were transformed into map distances according to the Kosambi function [22]. Subsequently, to obtain a common linkage map among the populations, smoothing was performed among the three populations using the ‘loess’ function in the ‘stats’ package version 4.1.2 in R.

Based on the common linkage map described above, QTL analysis was performed for each population using the ‘cim’ function in the ‘qtl’ package version 1.60 in R [23]. The window size was set to 10 cM, and the covariate markers were set to five. The logarithm of odds (LOD) scores were determined using 10,000 permutations. The targeted traits in the QTL analysis were PA, RA, and ANT in the F_3_ generation and PA, RA, yield, and flowering date in the F_4_ generation.

### 2.5. Genomic Heritability and GP Model

Genomic heritability was estimated for all traits described in Subsection 2.2, and single-and multi-trait GP models were constructed for each generation. GPs were made for each population and for three combined populations.

To evaluate GP accuracy, 10-fold cross-validation was performed and repeated 10 times. For each cross validation, the Pearson correlation between the observed and predicted values was calculated, and the average of these correlations was used as the prediction accuracy.

### 2.5.1. Single Trait GP Model

GBLUP [24] and BayesB [7] models were used for single-trait GP. The GBLUP model can be expressed as

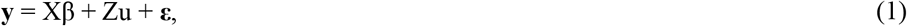

where n is the number of individuals, g is the number of genotypes, **y** is an n × 1 vector representing phenotypic values of a target trait, **Xβ** is an n × 1 vector corresponding to fixed effects, **Zu** is an n × 1 vector corresponding the term of random effects. In the prediction model for each population, **Xβ** represents an intercept, whereas in the model for all three populations, **X** represents an n × 3 design matrix, **β** is a 3 × 1 vector, and **Xβ** is a term correcting for the average effect of each population. The random effect **u** is a g × 1 vector corresponding to genotypic values that follow a multivariate normal distribution:

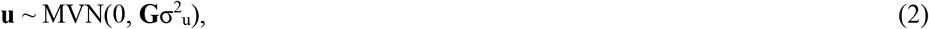

where **G** is a g × g additive genomic relationship matrix, and σ^2^_u_ is a genetic variance. In this study, the additive genomic relationship matrix **G** was computed based on the marker genotype data using the ‘calcGRM’ function in the ‘RAINBOWR’ package version 0.1.29 in R [25]. The additive relationship matrix G can be computed as a linear kernel of the marker genotype scaled by the allele frequency divided by the normalization constant, as described previously [26]. **ε** is an n × 1 error vector that follows a multivariate normal distribution;

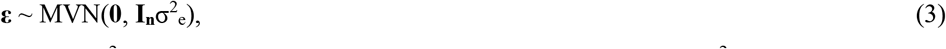

where **I**_**n**_ is an n × n identity matrix, and σ^2^_e_ is an error variance. We estimated the genetic variance σ^2^_u_ and the error variance σ^2^_e_ using the ‘EMM.cpp’ function in the ‘RAINBOWR’ package version 0.1.29 in R [24]. Based on the estimated genetic and error variances, the genomic heritability for each trait was calculated as h^2^ = σ^2^_u_ / (σ^2^_u_ + σ^2^_e_).

The BayesB model can be written as:

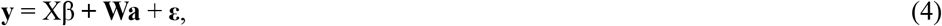

where m is the number of markers, **W** is an n × m matrix of the marker genotype, and **a** is an m × 1 vector representing the marker effects. The model was implemented using the ‘BGLR’ function in the ‘BGLR’ package version 1.1.0 in R [27].

### 2.5.2. Multi Trait GP Model

GBLUP [28] and BayesCπ [29] models were used for multi-trait GP. The GBLUP model is similar to Equation 1, but with the vector of phenotypic values **y** being extended to an nt × 1 vector through vertical alignment of the phenotypic values of the t traits. Additionally, the random effect **u** now follows a multivariate normal distribution **u** ∼ MVN(**0, K** ⨂ **G**), where **K** is a t × t genetic variance-covariance matrix between the t traits, and **G** is the g × g additive genetic matrix. The non-diagonal elements of **K** represent the covariances between traits. The error **ε** follows a multivariate normal distribution **ε** ∼ MVN(**0, R** ⨂ **I**_**n**_) where **R** is the t × t residual variance-covariance matrix of the traits, and **I**_**n**_ is the n × n identity matrix. Here, **A** ⨂ **B** represents the Kronecker product between the two matrices A and B. The BayesCπ model is similar to Equation 4. The model was also extended to that for multi-trait GP by considering the covariances between the different t traits at each marker. Furthermore, in the BayesB for the single-trait model, the prior distribution of each marker effect is a scaled-t distribution, whereas in the BayesCπ for the multi-trait model, a t × 1 vector of each marker effect for t traits follows a normal prior distribution. The multi-trait GP models were implemented using the ‘Multitrait’ function in ‘BGLR’ package version 1.1.0 in R [27].

## 3. Results

### 3.1. QTL Analysis

We performed QTL analysis in each population and each generation. In the F_3_ generation, the three target traits (PA, RA, and ANT) were investigated. In the F_4_ generation, the PA, RA, yield, and flowering date were investigated. Following composite interval mapping, 136 QTLs in the F_3_ generation and 268 QTLs in the F_4_ generation were detected above the threshold LOD scores. The number of QTLs consistent across multiple populations or two generations was 48 for PA, none for RA (Figure S1), two for ANT (Figure S2), two for yield (Figure S3), and 11 for flowering date (Figure S4). Among these QTLs, two QTLs for ANT and two for yield were detected at identical locations. These two QTLs may represent a single pleiotropic QTL. As shown in Figure 2a, when the genotype at the detected QTL was homozygous for ‘SekihoS8’ (red perilla), the phenotypic value of ANT content was also higher. In S844, no QTLs were detected, and all individuals were homozygous for ‘SekihoS8’ at the corresponding markers. However, when the genotype at the QTLs detected for yield was homozygous for ‘SekihoS8,’ the phenotypic value of yield was lower than that of the other genotypes (Figure 2b).

**Figure 2.**
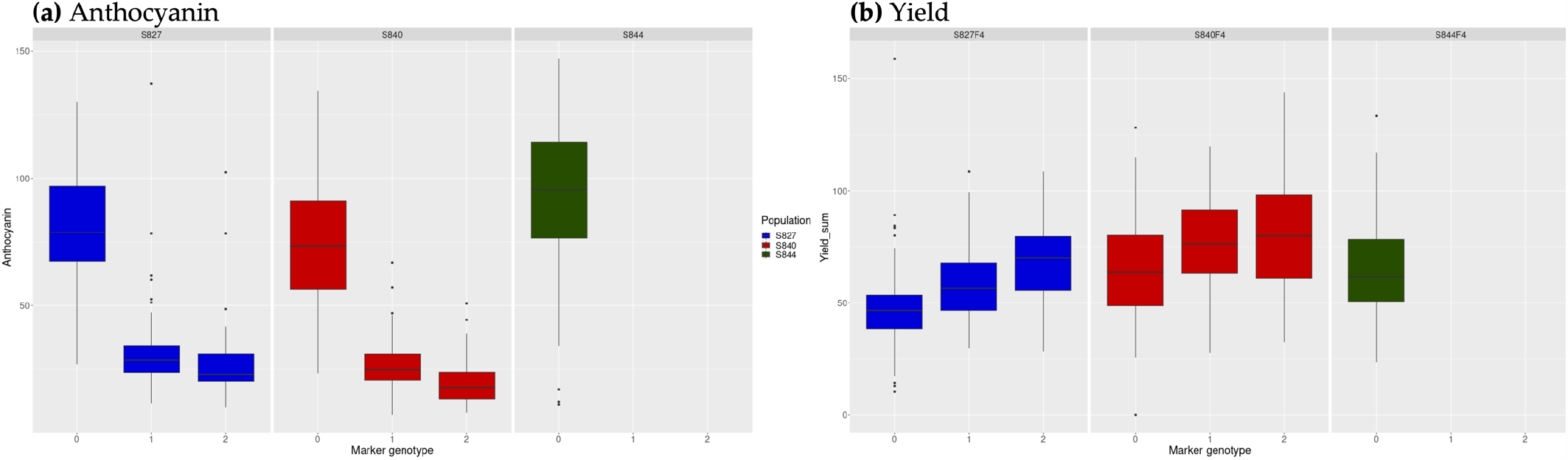
Genotypes at the detected quantitative trait loci (QTLs) identical to anthocyanin and yield. Genotype scores 0, 1, and 2 represent homozygous for ‘SekihoS8’, heterozygous, and homozygous for the other cross parents, respectively. Blue: S827; Red: S840; Green: S844. **(a)** The genotypes at the detected QTL on chr8 for anthocyanin (1^st^ measurement); **(b)** Genotypes at the detected QTL on chr8 for yield.

Eleven QTLs for flowering date were detected across multiple populations, of which one QTL was common to S827 and S844 and 10 QTLs were common to S840 and S844. We found that when the genotype at the detected QTLs was homozygous for ‘SekihoS8,’ the plant had an earlier flowering date (Figure S5).

Among the QTLs detected in PA, 16 were common to S827 and S844 on chr5. Twenty-eight QTLs were detected in S840 on chr5 and chr7, which were common to the F_3_ and F_4_ generations, and were at different positions from the QTLs detected in the other two populations (Figure 3). For QTLs commonly detected in S827 and S844, the plants with genotypes homozygous for ‘SekihoS8’ had lower PA contents (Figure S6a). In contrast, for QTLs detected only in S840, plants with genotypes homozygous for ‘SekihoS8’ had high PA contents, whereas those with genotypes homozygous for ‘st40’ had a PA content close to zero (Figure S6b).

**Figure 3.**
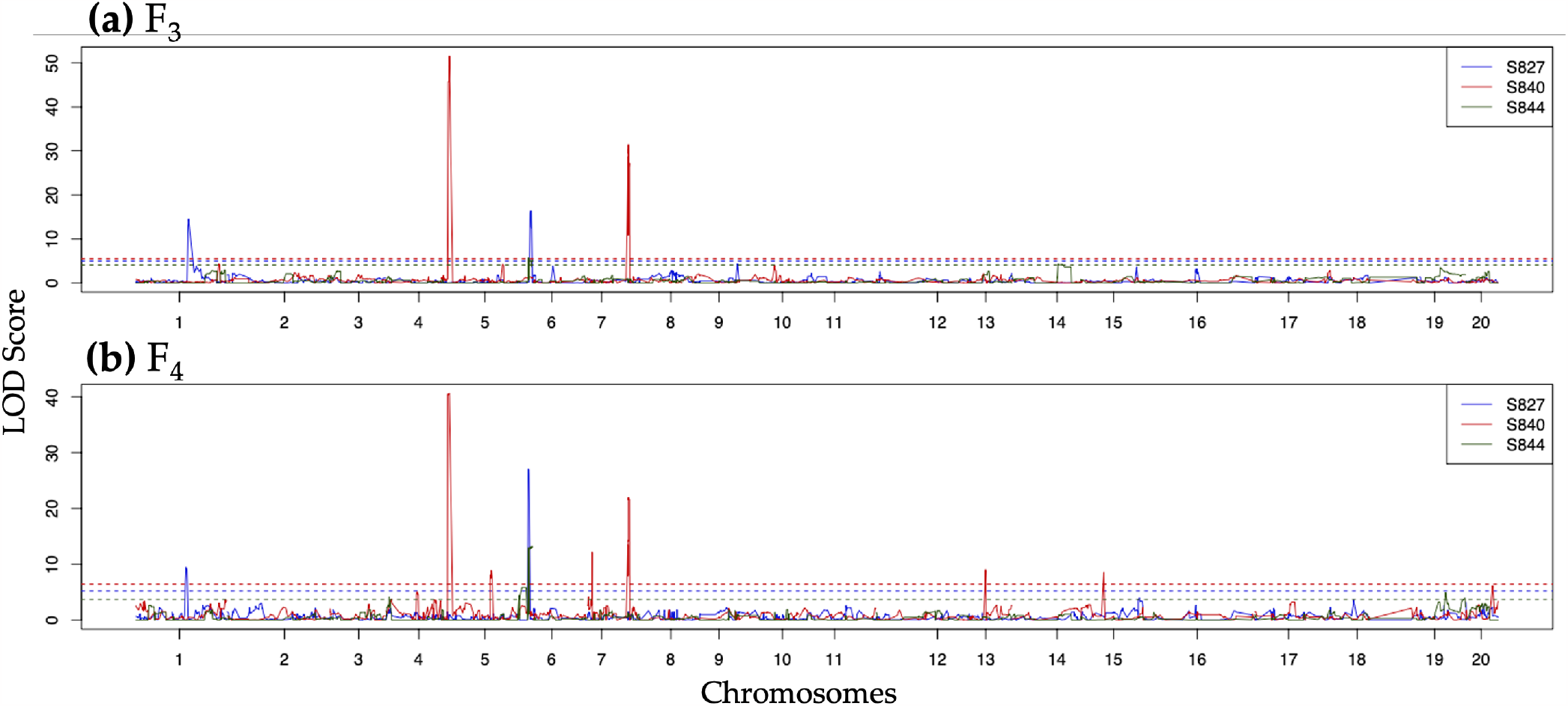
Quantitative trait loci (QTLs) detected for perillaldehyde in S827 (blue), S840 (red), and S844 (green). **(a)** QTLs detected in the F_3_ population; **(b)** QTLs detected in the F_4_ population. Dashed lines represent the logarithm of odds (LOD) threshold estimated by 10,000 permutations for each population.

### 3.2. Genomic Heritability

The genomic heritability of all traits mentioned in Subsection 2.2 estimated using each population and the three populations combined is shown in Tables 1 and 2. Genomic heritability varied substantially among traits and populations. Among all the traits, RA had the lowest genomic heritability, with values ranging from 0.151 to 0.484. In contrast, PA, ANT, yield, and flowering dates showed considerably higher genomic heritability. Among the three populations, the genomic heritability estimated using S844 was consistently the lowest for almost all traits, except yield. When the genomic heritability was estimated using the three populations combined, it was always greater than the average genomic heritability estimated using each population. For the flowering date, the genomic heritability estimated using the three populations combined was 0.879, which was the highest among all the estimates made using each population.

**Table 1.**
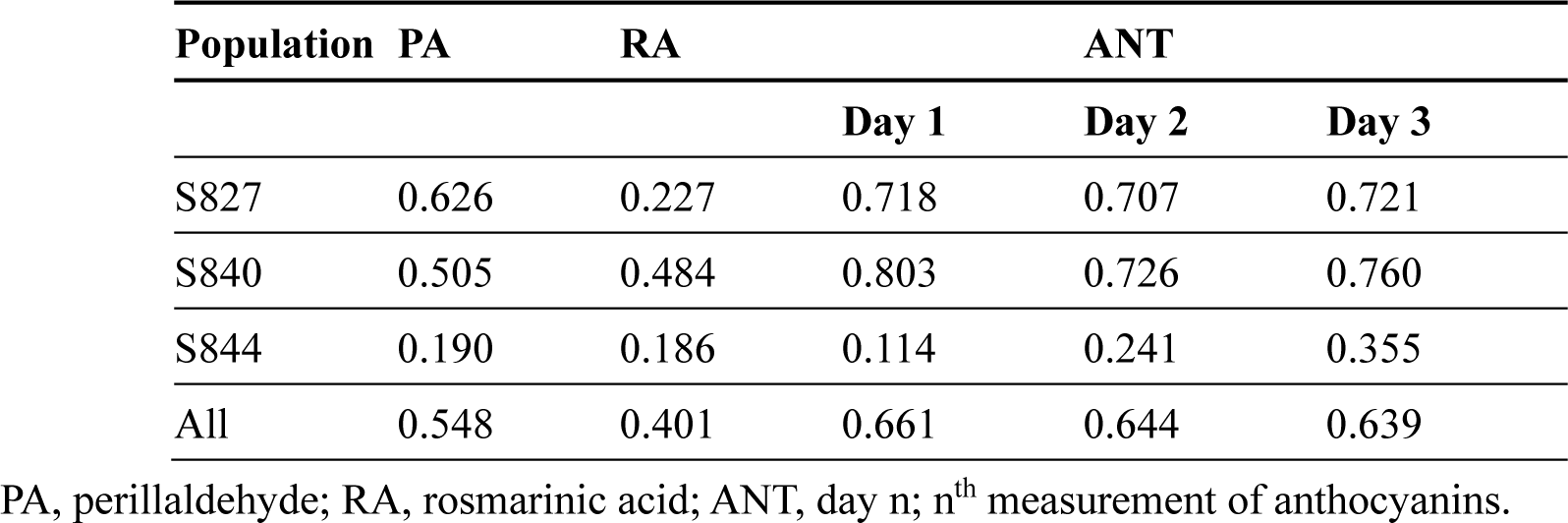
Estimated genomic heritability in F_3_ generation using each population and three populations combined.

**Table 2.** Estimated genomic heritability in F_4_ generation using each population and three populations combined.

| Population | PA | RA | YI | FD |
| --- | --- | --- | --- | --- |
| S827 | 0.766 | 0.394 | 0.634 | 0.869 |
| S840 | 0.796 | 0.389 | 0.605 | 0.848 |
| S844 | 0.449 | 0.151 | 0.715 | 0.691 |
| All | 0.735 | 0.350 | 0.702 | 0.879 |

PA, perillaldehyde; RA, rosmarinic acid; YI, yield; FD, flowering date.

### 3.3. Genetic Correlation

The multi-trait GBLUP model described in Subsection 2.5.2 was used to estimate genetic correlations among all traits. As shown in Figure 4, variations were observed in the positivity, negativity, and magnitude of genetic correlations among different generations and populations; however, for ANT, the correlations were positively strong across the three measurements in all populations. For genetic correlations estimated using three populations combined, two prime medicinal compounds, PA and RA, showed weak negative correlations with ANT (from -0.2 to -0.12), while PA and yield showed a moderate positive correlation. The genetic correlation between these two traits became more negative from the F_3_ to the F_4_ generation, except for S840. When genetic correlations were estimated for each population, S840 had the lowest absolute value for almost all traits.

**Figure 4.**
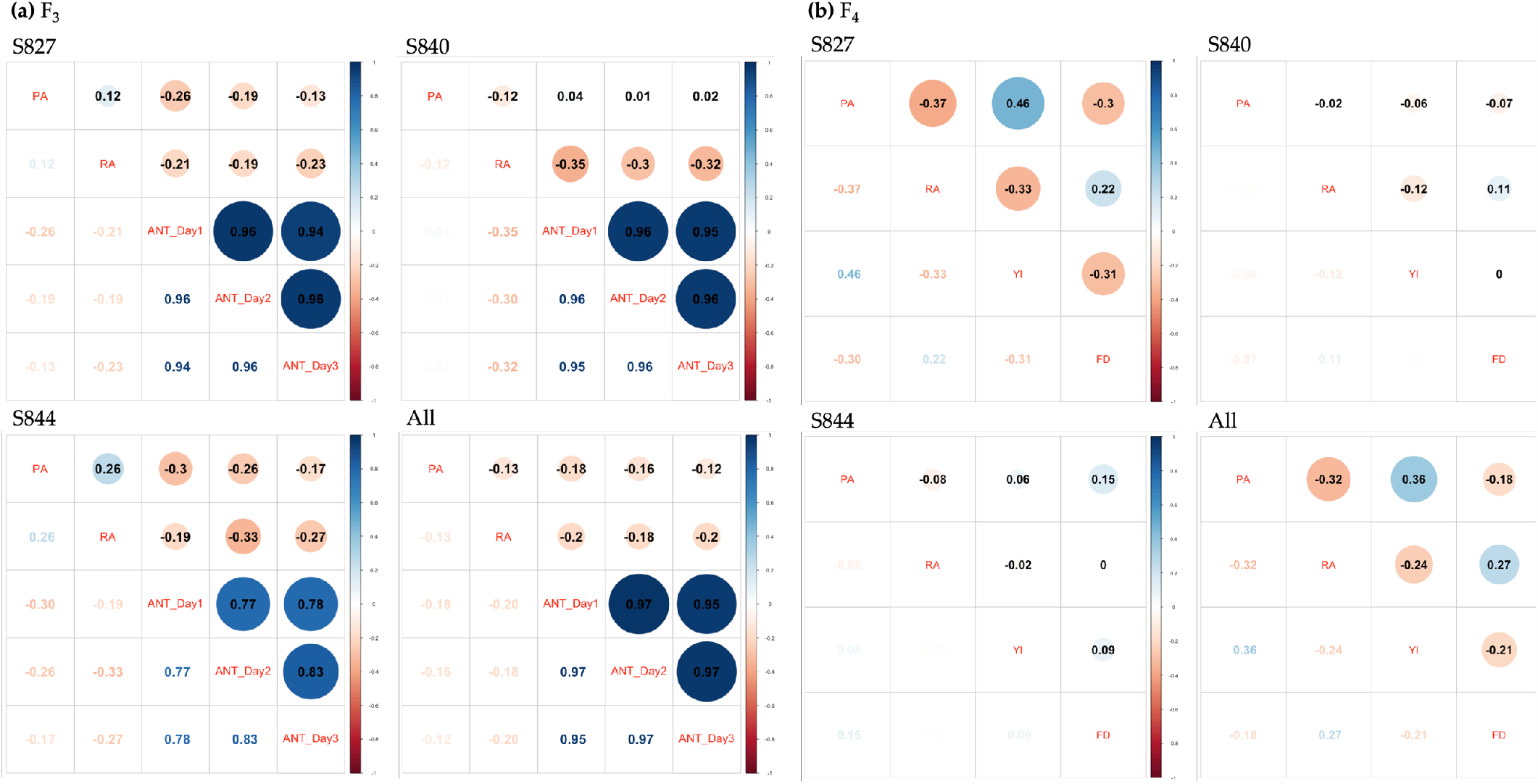
Genetic correlation among traits using each population and three populations combined. **(a)** Genetic correlations using the F_3_ population; **(b)** Genetic correlations using the F_4_ population. PA: perillaldehyde, RA: rosmarinic acid, ANT_Dayn: n^th^ measurement of anthocyanin, YI: yield, FD: flowering date.

### 3.4. GP Accuracy

GP models were constructed for each F_3_ and F_4_ generation, using each population and three populations combined. The prediction accuracies of the two single-trait models (GBLUP and Bayes B) are shown in Figure 5. Traits and populations with low genomic heritability tended to have lower GP accuracy. When the models were constructed using each population, regardless of the GP model, S844 had the lowest accuracy for almost all traits; S827 had the highest accuracy for yield and flowering date; and S840 had the highest accuracy for PA, RA, and ANT. The models constructed using the three populations combined showed improved accuracies compared to models constructed using each population for PA and flowering date. When comparing the accuracies of the GBLUP and BayesB models, the BayesB model had better prediction accuracies for most traits, with values ranging from 0.0466% to 41.7% higher than those of GBLUP. However, the GBLUP model was superior for some traits and populations, particularly RA, with a 0.0547–4.25% higher accuracy than the BayesB model. Furthermore, the same GP model showed higher prediction accuracy when using the F_4_ population than the F_3_ population. When the multi-trait models (GBLUP and BayesCπ) were used, the prediction accuracies obtained were similar to those obtained using the single-trait model (Figure S7).

**Figure 5.**
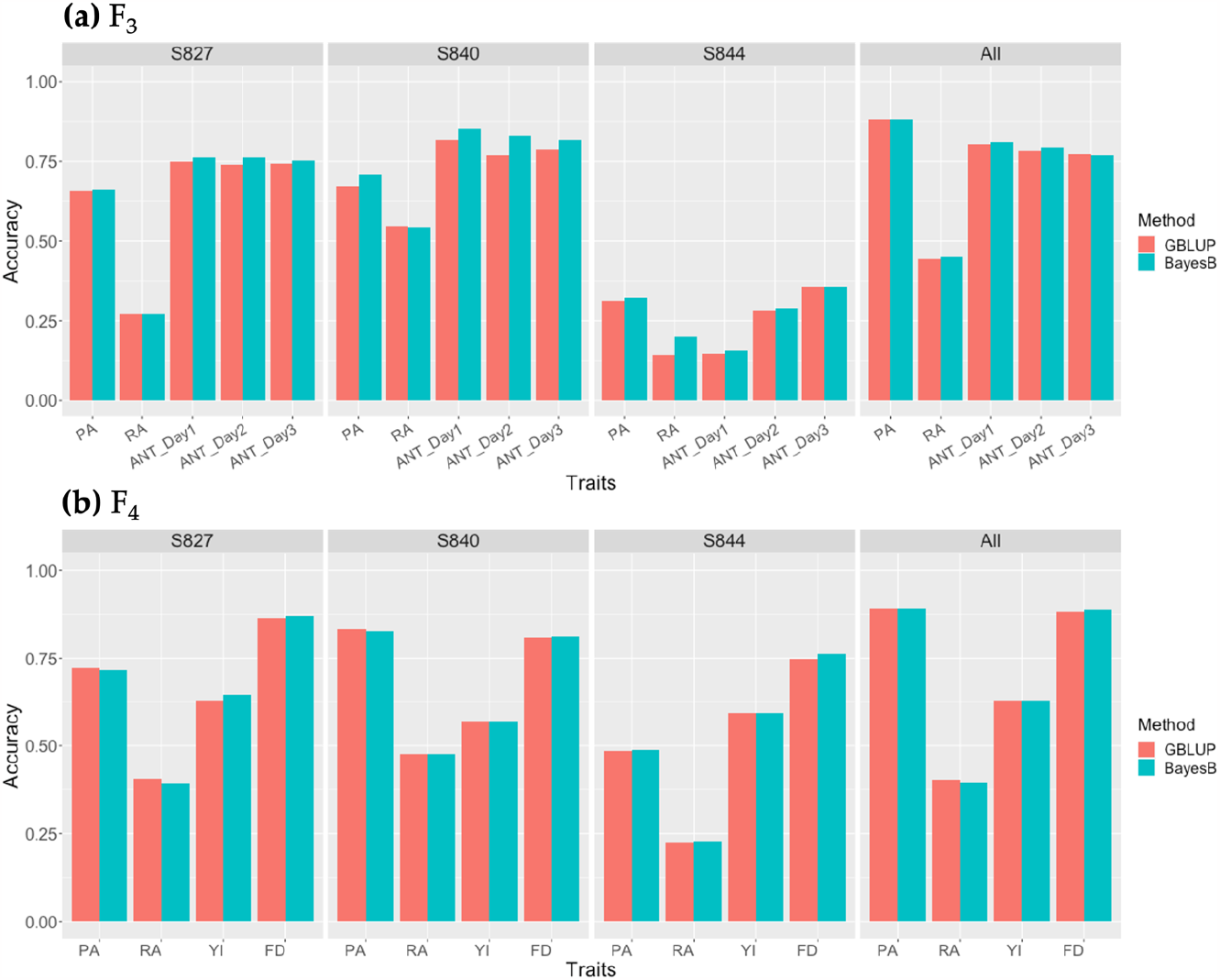
Prediction accuracy of single-trait genomic prediction (GP) by the GBLUP (red) and BayesB (cyan) models using each population and three populations combined. **(a)** Prediction accuracy using the F_3_ population; **(b)** Prediction accuracy using the F_4_ population. PA: perillaldehyde, RA: rosmarinic acid, ANT_Dayn: n^th^ measurement of anthocyanin, YI: yield, FD: flowering date.

## 4. Discussion

In this study, we conducted QTL analysis and GP for the major medicinal compounds and other agronomic traits of *P. frutescens* to reveal the genetic mechanisms underlying these traits and to evaluate the efficiency of GS and MAS. The results of genetic correlations, heritability, and GP accuracy revealed that each population had different characteristics; however, reasonable values were obtained when the three populations were used together. The results of the QTL analysis for each trait showed that traits such as PA, ANT, and flowering date had significant QTLs with high LOD scores; however, no significant QTLs were detected in RA. These results revealed the genetic characteristics of each trait. In Subsection 3.1, examination of the association between marker genotypes at the QTL and phenotypes of ANT revealed that plants with the ‘SekihoS8’ homozygous genotype had a significantly higher ANT content. ‘SekihoS8’ is a red perilla variety, and red perilla is reported to accumulate more ANT in the leaves [2, 3]. In contrast, the green perilla line, the other cross-parent, has been reported to accumulate minimal amounts of ANT in the leaves. Therefore, the detected QTLs were considered important gene loci associated with the ANT metabolic pathway. Zhang et al. performed a Genome-Wide Association Study for ANT and detected a strong signal on chr8, which was assumed to be the MYB transcription factor gene that regulates ANT biosynthesis in vegetative plant tissues [30]. The two QTLs for ANT detected in our study were also located on chr8 and had extremely high LOD scores, which is consistent with the results of Zhang et al. [30]. QTLs for yield were identical to those for ANT, but yield tended to be lower when the genotype at these QTLs was homozygous of the ‘SekihoS8’ allele. The QTLs for ANT had the opposite effect on yield; however, simultaneous enhancement of both traits during breeding was not difficult, as even though these QTLs have notable effects on ANT, yield is also substantially affected by other markers. Therefore, individuals with high ANT content can be selected by MAS with these markers, and an improvement in yield can be achieved by combining the effects of other markers with GS. For PA, the QTLs detected only in S840 were at positions different from those detected in S827 and S844. When the marker genotypes of the QTLs detected only in S840 were homozygous for st40, the PA content was nearly zero. PA is an essential oil that produces a strong odor. During cultivation, st40 smelled considerably different from st27 and st44. These results suggest that PA production by the plant is controlled by the QTLs located on chr5 and chr7 in S840. Furthermore, among the perilla plants that produce PA, the QTLs that determine the amount of accumulated PA may be located at different positions on chr5, which were detected in S827 and S844. Among the detected QTLs, one QTL was common to both S827 and S844, and 10 QTLs were common to both S840 and S844. Kang et al. performed a QTL analysis on traits related to flowering time and detected six QTLs that were identified as perilla orthologs that regulate flowering time [31]. Because the linkage groups in their study did not converge to 20, the number of chromosomes in perilla, we cannot confirm whether this is consistent with the QTLs we detected.

According to the estimation results of genomic heritability and GP, all traits except for RA had considerably high genomic heritability, indicating that improvement through GS was effective for these traits. Even for RA, which had the lowest genomic heritability among all traits, the genomic heritability was approximately 0.4 in S827 and S840. Because we evaluated the genomic heritability of each trait based only on additive genetic effects in this study, selection accuracy could be further improved by considering non-additive genetic effects, such as dominance and epistatic effects, in the GP model, even for traits with low genomic heritability, such as RA [32].

In terms of the comparison of prediction accuracy between different GP models, the BayesB and GBLUP models differed depending on the traits. This is because the fitness of either model depends on whether each trait is controlled by a small number of QTLs with large effects, or by an accumulation of polygenes with small effects. For most traits, the BayesB model was more accurate than the GBLUP model; however, for RA, the GBLUP model was 4.2% more accurate than the BayesB model. This indicates that RA is regulated by a large number of genes. In contrast, PA, ANT, and the flowering date were controlled by a few genes with large effects. Such differences in the genetic mechanisms of the traits were also evident in the QTL analysis results.

The accuracy of genomic selection depends not only on genomic heritability but also on factors such as the strength of linkage disequilibrium (LD), number of QTLs, and population size [33]. For traits with low heritability, marker density may affect the accuracy of GP, with some studies showing that accuracy improved when the LD between adjacent markers was >0.2 [34]. Considering the relationship between marker density and GP prediction accuracy, S840 of the three populations showed the highest prediction accuracy for most traits, probably because 1,432 SNPs were used for prediction. This was approximately 2.5 times higher than that of the 579 SNPs used in S844, which had the lowest prediction accuracy among the three populations.

Furthermore, the evaluation of the multi-trait GP model, the prediction accuracy was similar to that of the single-trait GP model. One of the reasons for this may be that the genetic correlation among traits estimated in Subsection 3.3 was not strong. Another notable result was that the genetic correlations among traits differed for each population. During breeding, an absence of strong negative correlations between the target traits is highly desirable; however, according to the results in Subsection 3.3, some populations showed slightly negative genetic correlations for some traits. These negative genetic correlations can be reduced by simultaneously using multiple populations with different genetic backgrounds during breeding.

Finally, we examined the possibility of genomic breeding of red perilla as a medicinal plant and discussed the different genetic mechanisms underlying the regulation of the medicinal and other agronomic traits. Therefore, it is important to use different GP models and selection methods that match the genetic characteristics of each trait. Moreover, the results of this study indicate that simultaneously using multiple populations with different characteristics, such as different genetic correlations among target traits and different positions of QTLs, for breeding can result in multiple trait improvements when the size of each population is sufficient to build the prediction model. Our research contributes to the study of the genetic mechanisms underlying the production of medicinal compounds in perilla, as well as the acceleration of medicinal plant breeding using MAS and GS.

## Supplementary Materials

Figure S1: QTLs detected for rosmarinic acid in S827, S840, and S844. Figure S2: QTLs detected for anthocyanins in S827, S840, and S844. Figure S3. QTLs detected for yield in S827, S840, and S844. Figure S4: QTLs detected for flowering dates at S827, S840, and S844. Figure S5: Genotypes at the QTL detected on chr19 for flowering date. Figure S6: Genotypes of detected QTLs for perillaldehyde. Figure S7: Prediction accuracy of multi-trait GP by the GBLUP and BayesCπ models using each population and three populations combined.

## Author Contributions

Data curation, formal analysis, software, visualization, writing – original draft preparation, S. K.; conceptualization, K. S., K. H., T. T., M. S., and H. I.; formal analysis, K. H.; funding acquisition, T. T. and H. I.; investigation, T. T. and Y. A.; resources, T. T., M. S., T. K., K. S., and S. I.; writing—review and editing, supervision, project administration, and H. I. All authors have read and agreed to the published version of the manuscript.

## Institutional Review Board Statement

Not applicable.

## Informed Consent Statement

Not applicable.

## Data Availability Statement

All other relevant data are available upon reasonable request.

## Acknowledgments

The seeds of the cross-parents of the three populations used in this study were provided by the Genebank Project for Agricultural Biological Resources of the National Institute of Agrobiological Sciences (NIAS). We are also grateful to the technical staff of TSUMURA & CO., Kazusa DNA Research Institute, and Motoyuki Ishimori who played important roles in selecting the cross-parents of the populations used in this study. This work was supported by JST OPERA (grant number: JPMJOP1851).

