## Supplemental File for "Assessing the potential for genome-assisted breeding in red perilla using quantitative trait locus analysis and genomic prediction"

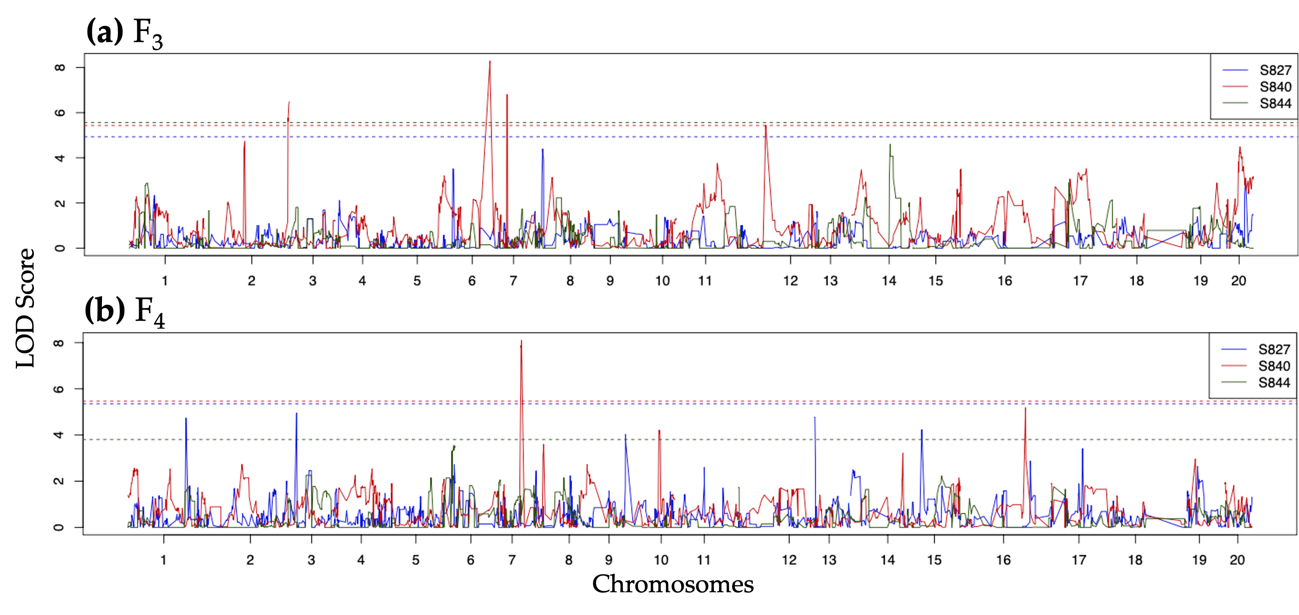

**Figure S1.** QTLs detected for rosmarinic acid in S827 (blue), S840 (red), and S844 (green). **(a)** QTLs detected in the F_3_ population; **(b)** QTLs detected. in the F_4_ population. Dashed lines represent the LOD threshold estimated by 10,000 permutations for each population.

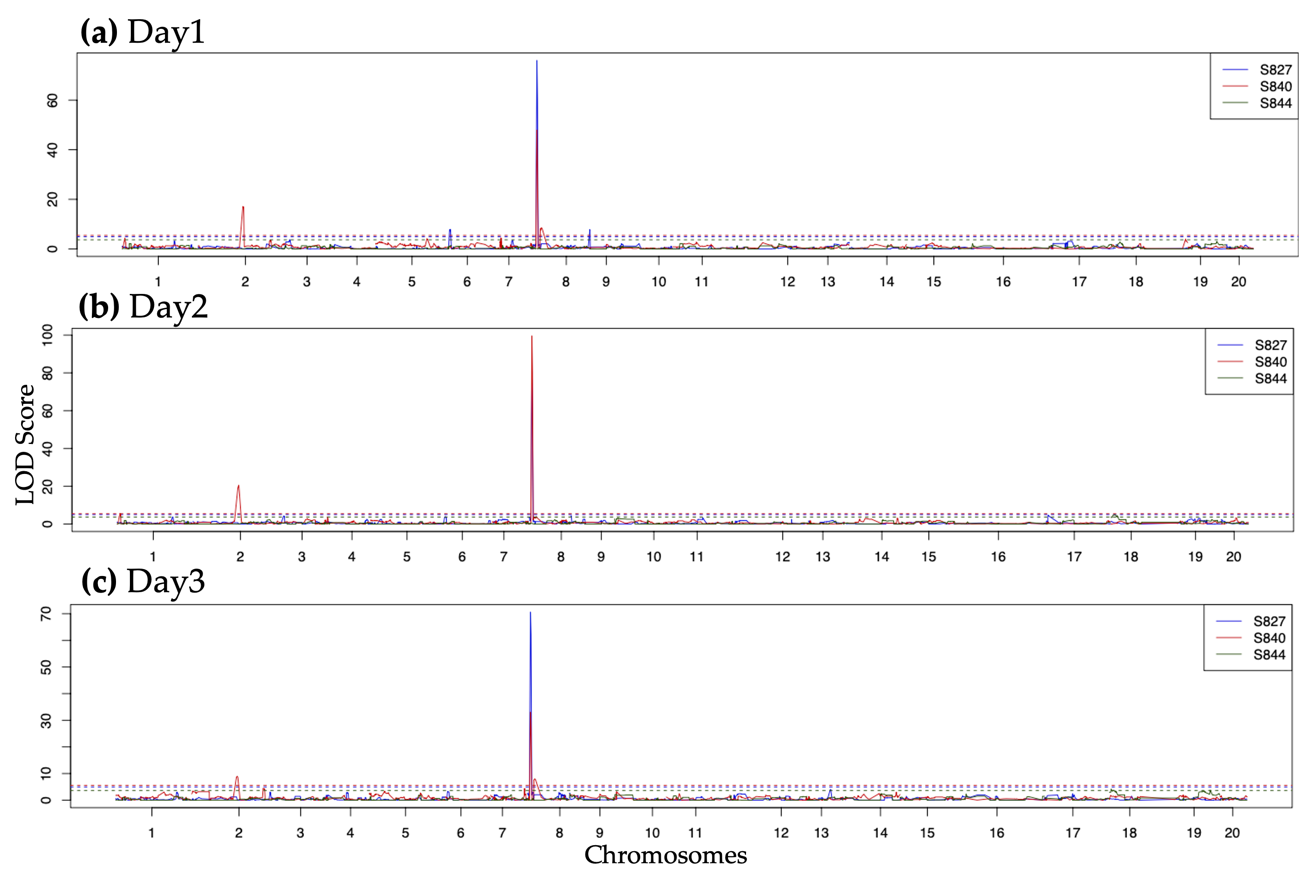

**Figure S2.** QTLs detected for anthocyanin in S827 (blue), S840 (red), and S844 (green). **(a)** QTLs detected in the 1^st^ measurement of anthocyanin; **(b)** QTLs detected in the 2^nd^ measurement of anthocyanin; **(c)** QTLs detected in the 3^rd^ measurement of anthocyanin. Dashed lines represent the LOD threshold estimated by 10,000 permutations for each population.

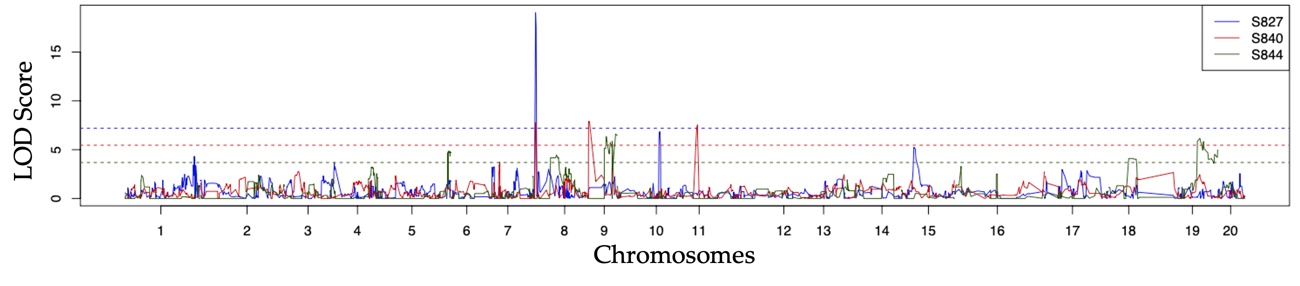

**Figure S3.** QTLs detected for yield in S827 (blue), S840 (red), and S844 (green). Dashed lines represent the LOD threshold estimated by 10,000 permutations for each population.

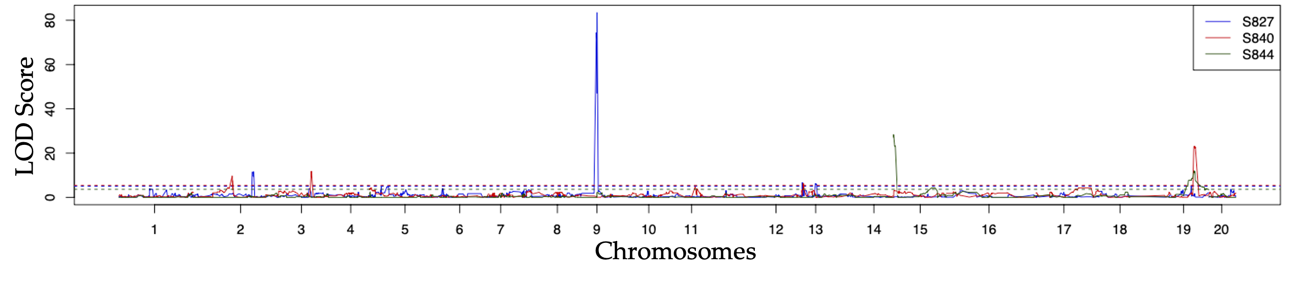

**Figure S4.** QTLs detected for flowering date in S827 (blue), S840 (red), and S844 (green). Dashed lines represent the LOD threshold estimated by 10,000 permutations for each population.

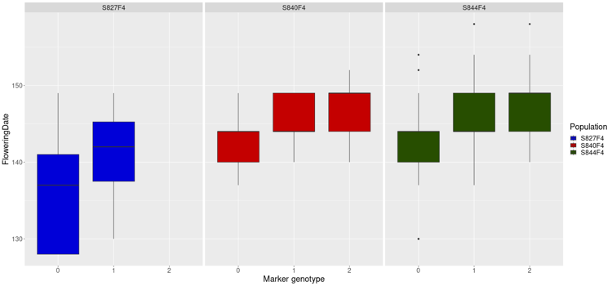

**Figure S5.** The genotypes at the detected QTL on chr19 for flowering date. Genotype scores 0, 1, and 2 represent homozygous for ‘SekihoS8’, heterozygous, and homozygous for the other cross parents, respectively. Blue: S827; Red: S840; Green: S844.

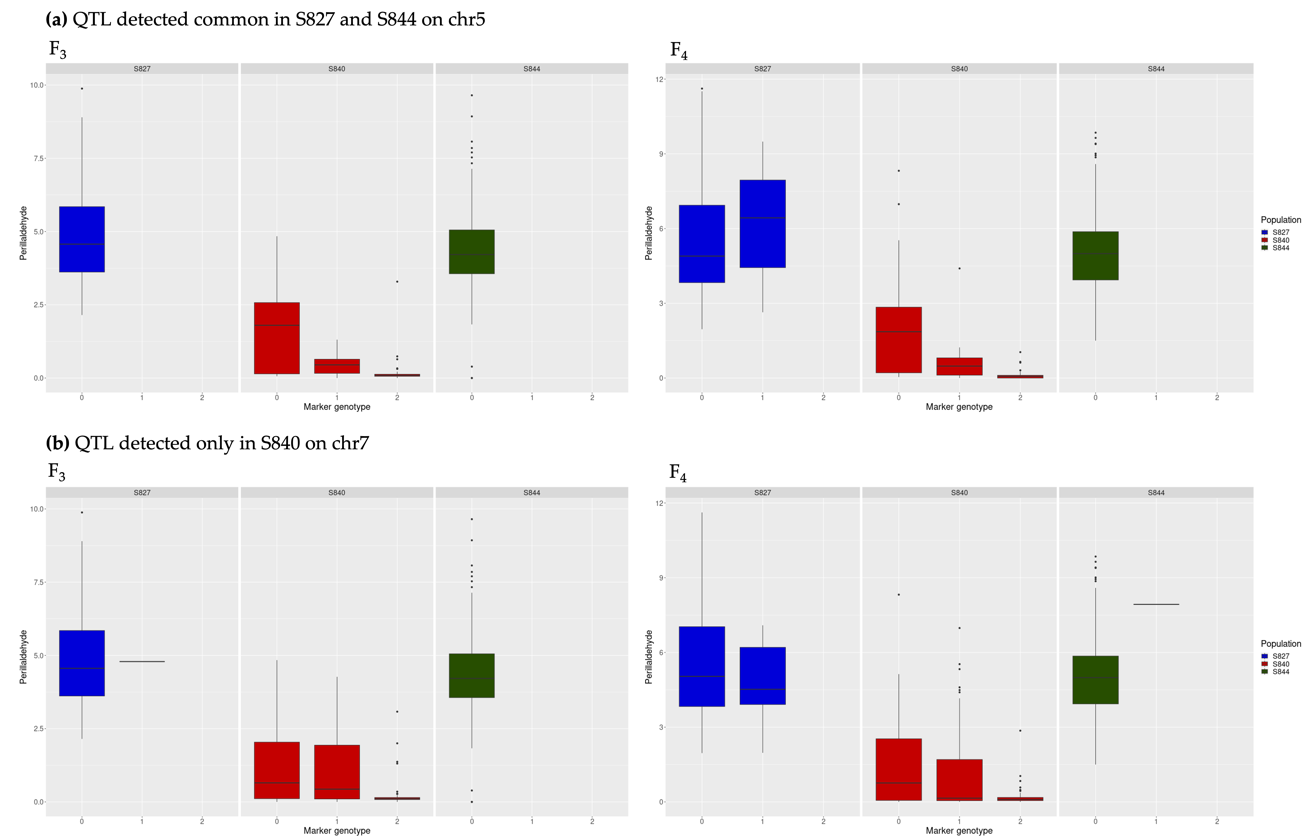

**Figure S6.** The genotypes at the detected QTLs for perillaldehyde. Genotype scores 0, 1, and 2 represent homozygous for ‘SekihoS8’, heterozygous, and homozygous for the other cross parents, respectively. The panel on the left side is for F_3_ generation and the panel on the right side is for F_4_ generation. Blue: S827; Red: S840; Green: S844. (**a**) The genotypes at the QTL detected in both S827 and S844 on chr5; (**b**) The genotypes at the QTL detected only in S840 on chr7.

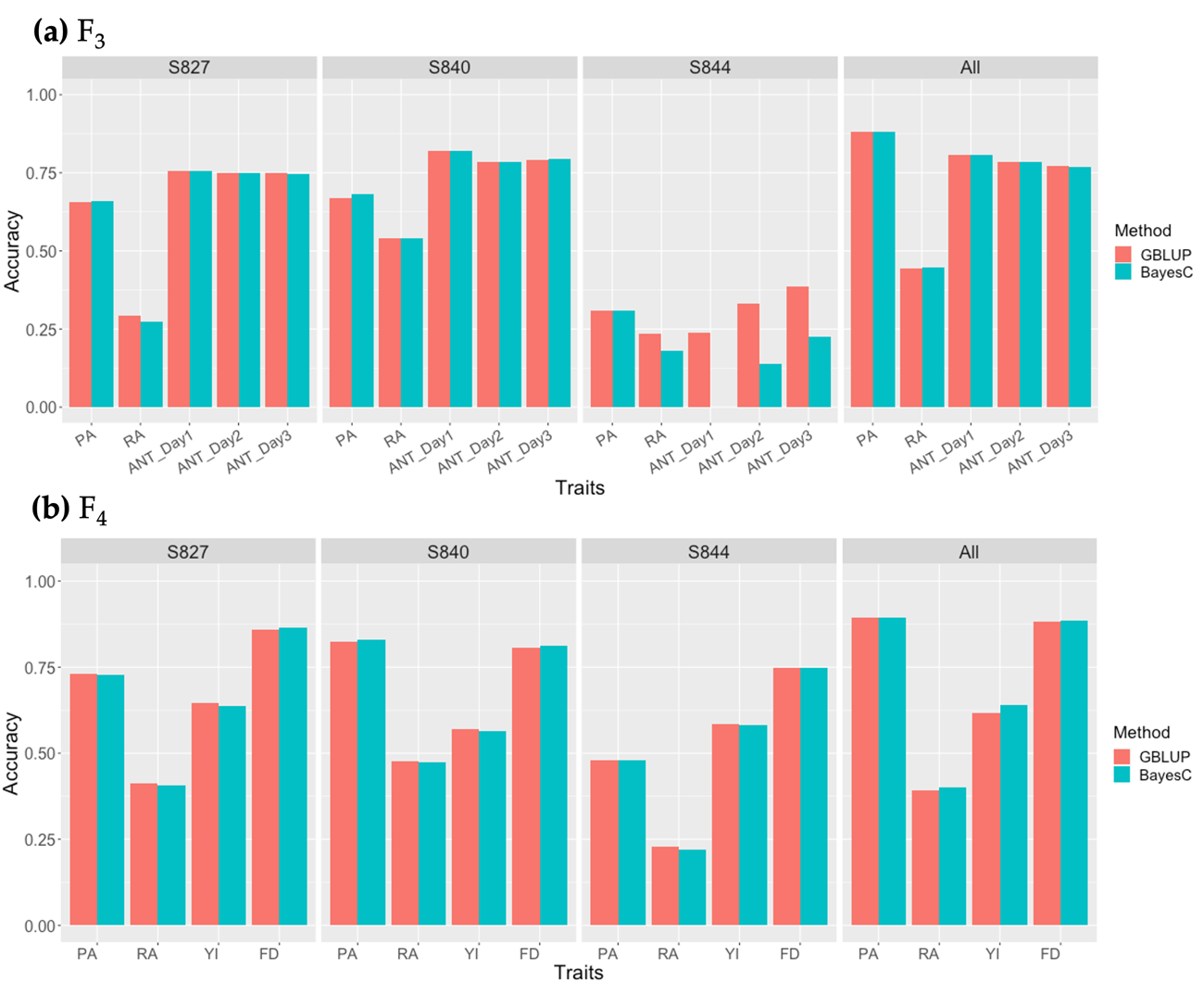

**Figure S7.** Prediction accuracy of multi-trait GP by the GBLUP (red) and BayesCπ (cyan) models using each population and three populations combined. **(a)** Prediction accuracy using the F_3_ population; **(b)** Prediction accuracy using the F_4_ population. PA: perillaldehyde, RA: rosmarinic acid, ANT_Dayn: n^th^ measurement of anthocyanin, YI: yield, FD: flowering date.
